# A natural killer cell gene signature predicts melanoma patient survival

**DOI:** 10.1101/375253

**Authors:** Joseph Cursons, Fernando Souza-Fonseca-Guimaraes, Ashley Anderson, Momeneh Foroutan, Soroor Hediyeh-Zadeh, Andreas Behren, Nicholas D. Huntington, Melissa J Davis

## Abstract

Animal models have demonstrated that natural killer (NK) cells can limit the metastatic dissemination of tumors, however their ability to combat established human tumors has been difficult to investigate.

A number of computational methods have been developed for the deconvolution of immune cell types within solid tumors. We have taken the NK cell gene signatures from several tools, then curated and expanded this list using recent reports from the literature. Using a gene set scoring method to investigate RNA-seq data from The Cancer Genome Atlas (TCGA) we show that patients with metastatic cutaneous melanoma have an improved survival rate if their tumor shows evidence of greater NK cell infiltration. Furthermore, these survival effects are enhanced in tumors which have a higher expression of NK cell stimuli such as IL-15, suggesting NK cells are part of a coordinated immune response within these patients. Using this signature we then examine transcriptomic data to identify tumor and stromal components which may influence the penetrance of NK cells into solid tumors.

These data support a role for NK cells in the regulation of human tumors and highlight potential survival effects associated with increased NK cell activity. Furthermore, our computational analysis identifies a number of potential targets which may help to unleash the anti-tumor potential of NK cells as we enter the age of immunotherapy.

## Introduction

Natural killer (NK) cells are an essential component of the innate immune system, playing a critical role in the clearance of cells which carry a viral burden, or have undergone oncogenic transformation. They are a subset of innate lymphoid cells with an exquisite cytotoxic ability, allowing them to effectively kill tumor cells even at a relatively low ratio (e.g. 1:1) [1]. It has recently been demonstrated that NK cells are important for stimulating the anti-tumor immune response, recruiting conventional type-1 dendritic cells (cDC1) through chemokine signaling (*via.* XCL-1 & CCL-5/RANTES; Fig. 1), ultimately resulting in the generation of a robust T-cell response [2]. The cytotoxic mechanisms used by NK cells share many similarities to CD8^+^ T-cells (Fig. 1), including secretion of granzymes (e.g. GZMA, GZMB, GZMK, GZMM) and perforin (PRF1) [3, 4].

A number of in vivo studies have demonstrated a role for NK cells in limiting the metastatic dissemination of melanoma [5-8] and there is growing interest in the targeting of NK cells for novel immunotherapeutics [9]. Important regulators of NK cell activity include the cytokines IL-15 [10], IL-12 [11] and IL-18 [12], chemokines such as CCL5 (RANTES) [13], growth factors such as TGF-β [14, 15], and the intracellular JAK-STAT signaling component CIS [16]. Evidence suggest that modulation of NK cell populations is feasible for cancer treatment [17], and treatments based upon systemic administration of IL-15 constructs have shown promise in leukemic and solid tumors [18-20]. Thus, while IL-15 provides a promising treatment to stimulate immune targeting of cancers, we note that cytokine-mediated NK cell activation and expansion can be increased in combination with IL-12 and IL-18 [21] or further amplified through deletion of CIS (encoded by *CISH*), an important negative regulator of cytokine signaling and effector function, such that *Cish*^-/-^ mice are resistant to a range of metastatic cancers [16]. Accordingly, we are yet to fully elucidate the range of molecular regulatory systems that control NK cell activity *in vivo.*

Advances in sequencing technology and associated methods for data analysis over recent decades have allowed the application of transcriptomic profiling to complex tumor samples [22, 23]. Resulting data have enabled the development of mathematical methods, such as CIBERSORT [24] that infer the relative abundance of immune cells which have infiltrated into solid tumor samples. While these tools have provided insight into the nature and scope of immune infiltration [25-28], there are further opportunities to capitalize on public tumor transcriptomic data in order to identify how changes in tumor phenotype are associated with changes in the relative abundance of immune sub-populations.

Phenotype-switching is an important regulatory program involved in the progression of melanoma [29] which has been linked to vemurafenib resistance [30] and general drug resistance [31]. It allows tumor cells to transition between proliferative (“epithelial-like”) and invasive (“mesenchymal-like”) behaviours, and in melanoma, there is strong evidence that TGF-β is an important driver of EMT/phenotype-switching programs [32], mediated in part by signaling molecules such as thrombospondin 1 [33]. This work examines the LM-MEL panel of melanoma cell lines [34] which we have extensively profiled in the context of phenotype switching [33, 35].

**Figure 1:**
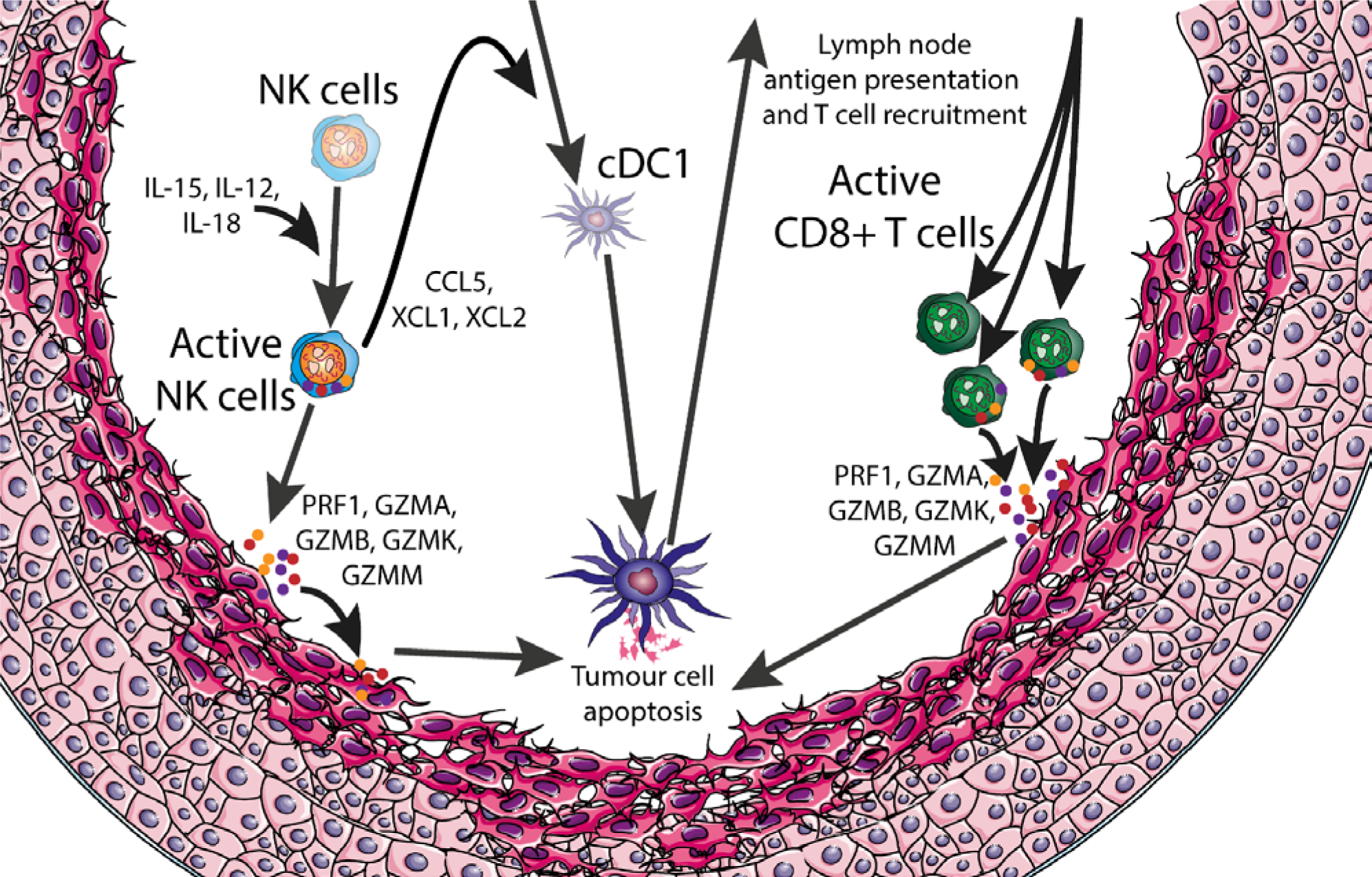
Immune targeting of tumours. A schematic overview of the intercellular signalling and microenvironment factors which can impact on anti-tumour immune responses. NK cells: natural killer cells; cDC1: conventional type-1 dendritic cells.

We have recently developed a single-sample gene set scoring method which uses a rank-based metric to quantify the relative enrichment of a gene set within a sample transcriptome [36]. We have combined NK cell signatures from the LM22/CIBERSORT [24] and LM7 [37] gene sets and curated this list to produce a gene set which reflects the relative abundance of NK cells within a tumor sample. As melanomas are highly-immunogenic, we have focused upon the analysis of TCGA skin cutaneous melanoma (SKCM) data [38] demonstrating that the relative expression of NK cell genes within metastatic tumors is associated with a strong survival advantage. Using the SKCM data we show how our scoring approach can be used to explore putative modulators of NK cell activity by examining their association with NK score and survival effects associated with their expression.

## Materials and Methods

### Data

**Table 1.** Data used in this report. Sources and references for the transcriptomic data, and patient data survival data for the TCGA samples.

| Resource | Data source & identifier | Reference |
| --- | --- | --- |
| TCGA SKCM | NIH Genomic Data Commons:<br>https://gdc.cancer.gov/ | [38] |
| RNA-seq data for sorted immune populations | NCBI Gene Expression Omnibus: GSE60424<br>https://www.ncbi.nlm.nih.gov/geo/ | [39] |
| Microarray data for sorted immune populations | NCBI Gene Expression Omnibus: GSE24759<br>https://www.ncbi.nlm.nih.gov/geo/ | [40] |
| LM-MEL melanoma cell line panel | EBI Array Express: E-MTAB-1496<br>https://www.ebi.ac.uk/arrayexpress/ | [34] |

Data used in this study are available from listed repositories (Table 1). For TCGA SKCM data, RSEM abundance data without normalization were downloaded directly from the genomic data commons. For sorted immune cell populations (GSE60424 & GSE24759) and melanoma cell line data (E-MTAB-1496), processed transcript abundances were downloaded and used directly. For GSE24759, only samples derived from peripheral blood were used, data from colony-forming samples was excluded to exclude culturing effects, and CD56-/CD16+/CD3-mature NK cell data were excluded due to apparent batch effects (Fig. S1). In cases with gene multi-mapping (multiple probes/probe sets per gene), median values were used.

As noted (Fig. S2A & B), there appear to be large survival differences between patients with primary and metastatic tumors. To avoid confounding effects from this, unless otherwise stated, we have focused on patients with metastatic tumors only who also had valid age and survival data. One patient with both a metastatic and primary sample was excluded.

### Computational Tools

The computational analysis was performed using python v3.6 together with pandas [41] for data handling, scipy [42] and numpy [43] for numerical calculations and matplotlib [44] for plotting. Gene set scoring was performed using the *singscore* gene set scoring approach [36].

### Relative log expression & principal component analysis

For relative log expression, log-transformed transcript abundance data (as downloaded directly from GEO) were median-centered for each gene, and then within each sample the difference between the observed and population median of each gene was calculated. For the principal component analysis, genes with an expression level above the 10^th^ percentile (5.34) within at least 4 samples (corresponding to the smallest sample group) were retained. Data were normalised using the sklearn [45] `StandardScaler` function, before calculating the principal components using the sklearn `PCA` function.

### Immune gene sets

A preliminary NK cell gene set was created from the CIBERSORT (LM22) active and resting NK cell gene sets [24] and the NK cell gene set from the LM7 gene sets [37]. These genes were further curated against a range of reports which have studied both human and mouse NK cell populations and the final gene list is given in Table S1. The TCGA immune gene set used for scoring was derived from Supplemental Table S4A and the TCGA classified ‘Immune high’ patients were taken from Supplemental Table 1D of the original SKCM manuscript [38].

### Gene set scoring

Gene set scoring was performed using the *singscore* approach [36]. Briefly, genes are ranked by increasing transcript abundance and for a set of target genes the mean rank is calculated and normalised against theoretical minimum and maximum values. If directional gene lists are provided (i.e. a set of genes expected to be upregulated and a set of genes to be downregulated), as with the TGF-β EMT signature [32] then the mean rank of expected up-regulated genes is calculated from genes ranked by increasing abundance, while the mean rank of expected down-regulated genes is calculated from genes ranked by decreasing abundance, and these values are then normalised and summed. Accordingly, a high gene set score indicates that the pattern of gene expression in a sample is concordant with the pattern captured by the gene expression signature.

### Survival analyses

All survival analyses, including the construction of Cox proportional hazard models and the generation of Kaplan-Meier survival curves were performed using the python package lifelines (v. 0.14.6; DOI: 10.5281/zenodo.1303381) with standard parameters.

### Code availability

All computational scripts used in this work will be made freely available from our GitHub repository:

https://github.com/DavisLaboratory/NK_scoring

An alternative implementation of this analysis using R/Bioconductor libraries has also been made available.

## Results

### Cutaneous melanoma is generally associated with a strong immunogenic response

Cutaneous melanoma is an ideal target for immunotherapy as the high mutational burden of this malignancy is associated with the generation of neo-antigens which can induce an immune response [46]. Several reports demonstrate that immune infiltration signatures provide a strong prognostic indicator in melanoma [47-49], including the TCGA skin and cutaneous melanoma (SKCM) study which demonstrated that this effect was independent of the underlying genomic subtype of the melanoma [38].

Due to significant survival differences between patients with a primary or metastatic tumor (Fig. S1A & B) this report focusses on the 365 patients with only metastatic tumor samples. There are significant survival effects associated with the patient’s age at diagnosis (Fig. S1C), and while female patients had better survival rates (Fig. S1D), these effects were not significant (Table 2; Cox proportional hazards model, *p*-value = 0.40).

**Table 2.**
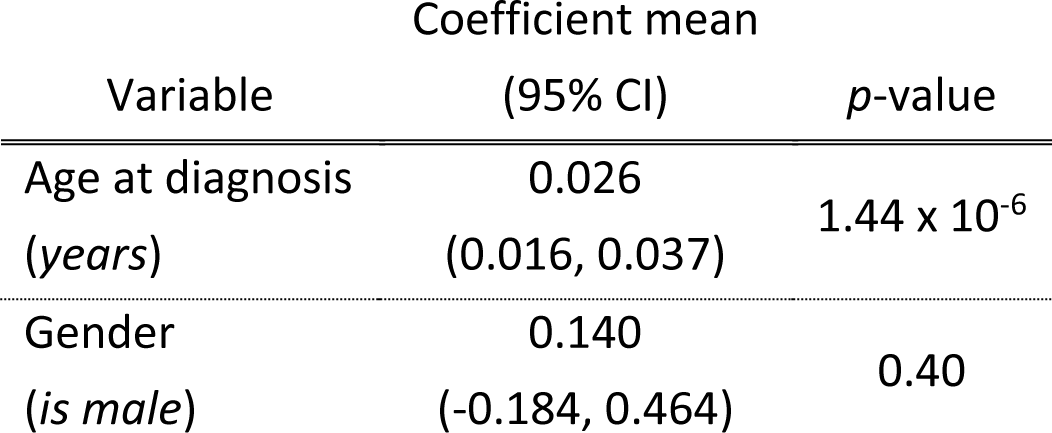
Covariate hazard coefficients for TCGA patients with metastatic melanoma. Variables were included within a Cox proportional hazards model and tested against the null hypothesis that the coefficient is equal to 0.

To analyze survival effects associated with individual genes we built a series of Cox proportional hazard models for each gene where patient age at diagnosis was included as the only covariate (together with transcript abundance for that gene). As shown in Fig. 2A, many genes with a hazard reduction have a known immune function (*genes in bold*); associated patient survival curves are shown for patient age (Fig. 2B) and a selection of genes (Fig. 2C-E). Higher expression of the hallmark inflammatory cytokine encoded by *IFNG* corresponds to improved survival outcomes (Fig. 2C), while a number of interferon-induced genes are further associated with a hazard reduction (*e.g. IRF1*, *IFITM1*; Fig. 2A). High tumor transcript abundances of the NK cell marker gene *KLRD1* (Fig. 2D; also known as *CD94*) or the cytokine IL-15 (Fig. 2E) which is an important regulator of NK cell [10, 50, 51] and T-cell activity [52, 53] are also associated with improved long-term survival outcomes. Finally, we note that transcript abundance for the *B2M* gene encoding beta-2 microglobulin has one of the most negative hazard coefficients, likely reflecting its role in MHC class I antigen presentation of neo-antigens to CD8 T cells and consistent with recent reports of the importance of this process for immune control of tumors [54]. Further, a truncation mutant of B2M can confer resistance to PD-1 blockade in melanoma [55], and mutations in B2M have been shown to disrupt immune surveillance in lung cancer [56]. The large negative hazard coefficient associated with *HAPLN3*, encoding a hyaluronan and proteoglycan link protein, suggests that this gene may warrant further investigation in the context of immune recognition and targeting.

**Figure 2:**
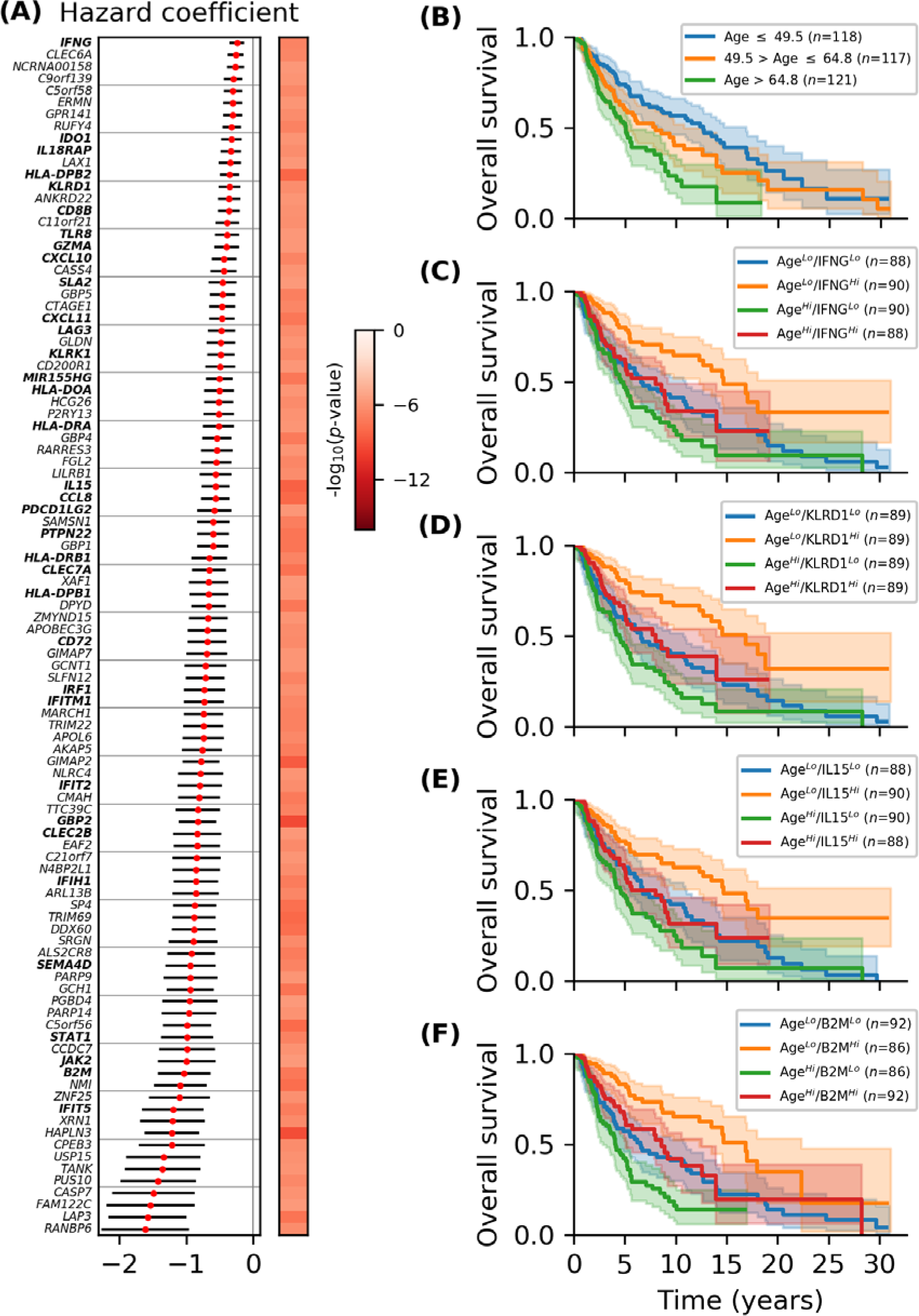
Hazard ratios associated with transcript abundance of individual genes. (**A**) A Cox proportional hazard model was created for each gene with patient age as the only co-variate. The top 50 genes were selected by significance and ranked by hazard coefficient (red dot, 95% confidence intervals shown with black lines). (**B**) Kaplan-Meier survival curves for patients with metastatic melanoma partitioned by age at diagnosis. (**C** - **F**) Kaplan-Meier survival curves for patients partitioned by age and (C) *IFNG*, (D) *KLRD1*, (E) *IL15*, or (F) *B2M* transcript abundance.

### Gene set scoring allows dimensional reduction of RNA-seq data

Gene set enrichment analyses are commonly used after differential expression to assess whether genes with the most significant changes are enriched for classifications of specific pathways or processes. An alternative ‘relative approach’ [23] is to analyze the gene expression patterns (transcript abundances) of individual samples and calculate the relative concordance of each one against specific gene set.

We have recently developed a gene set scoring approach [36] which uses the normalised mean rank of genes that are associated with a specific molecular phenotype or cellular behavior [32, 57]. With this approach, a difference in score between two samples can be related to the percentile change in mean rank of the gene set, providing a metric which summarizes the concordance between the gene expression profile of an individual sample and the specified gene sets. Using this scoring method with “Immune cluster” genes from the original TCGA SKCM publication [38] we can largely recapitulate the original sample clustering (Fig. S3), and furthermore, we can easily extend this analysis to samples that have more recently been added to the TCGA SCKM cohort.

### Developing a more specific transcriptomic signature for natural killer cells

A number of transcriptomic data deconvolution methods have generated gene signatures that are predictive of infiltration for specific immune cell sub-populations. Of note for this work, we examined the training data (transcriptomics from sorted immune cell populations) and NK cell signatures from the LM22 [24] and LM7 [37] gene sets. A common critique of immune deconvolution methods is the high co-linearity/cross-correlation between different signatures [37]. While this can largely be attributed to ‘marker genes’ which are common between immune cell subsets (demonstrated by similar positions of sorted cell populations in Fig. S1B & S1C), to an extent it also represents the cascading series of intercellular interactions that mediate immune activation within complex tissue samples. Accordingly, several immune-associated gene subsets are cross-correlated to a varying extent (Fig. S4). We combined the NK cell signatures from the LM7 and LM22 gene lists and curated this gene list to remove genes which are associated with a wide range of immune cell sub-populations (e.g. *IFNG* and *IL15*). The resulting 40 genes were used to score metastatic tumor samples from the TCGA SKCM cohort. The transcript abundance of each gene is shown for metastatic tumor samples sorted by their overall NK score (Fig. 3, at left), together with RNA-seq (at center) and microarray data (at right) for sorted immune cell populations. As shown (center panel) many of the NK cell marker genes are shared with CD4^+^ and CD8^+^ T-cell populations, however the resulting score is still greater within NK cell samples (for reference, a similar figure with the T-cell signature is given in Fig. S5). Of note, *CD244*, *GZMB*, *NKG7*, *XCL1* and *XCL2* all appear to have greater expression within NK cells relative to the T-cell sub-populations examined. Examining the microarray data from a much larger subset of sorted immune cell populations (Fig. 3, at right), NK cells have the highest resulting score. Again, there appears to be a subset of granulocyte and CD8^+^ T-cell samples which have relatively high NK scores, however these still tend to be lower than scores for the sorted NK cell populations.

**Figure 3:**
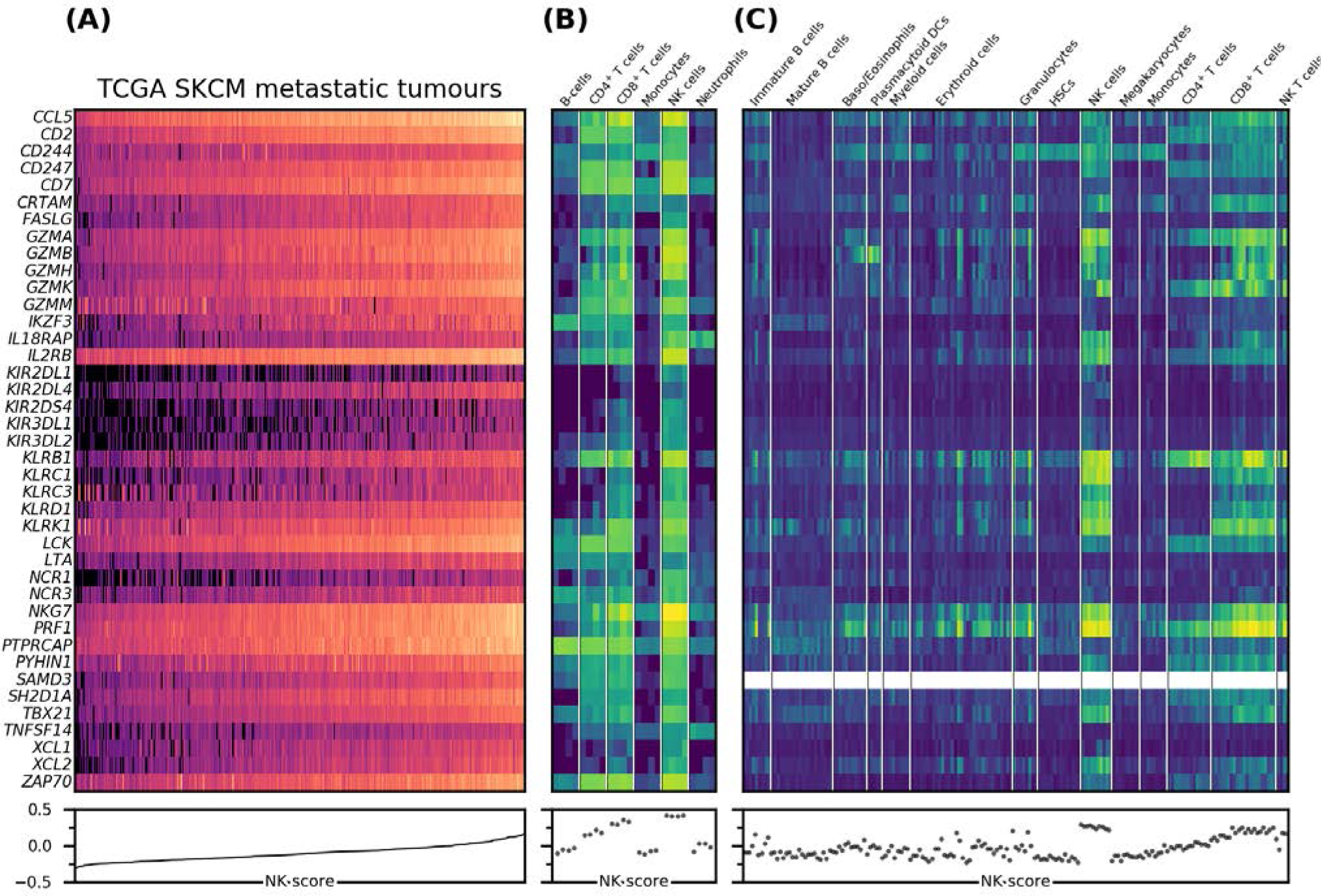
A refined natural killer cell gene signature in selected transcriptomic data sets. Transcript abundance of NK cell marker genes across (**A**) TCGA SKCM metastatic tumours ranked by NK score; (**B**) RNA-seq data for sorted immune cell populations (GSE60424); (**C**) microarray data for sorted immune cell populations (GSE24759). NK: natural killer; DCs: dendritic cells; HSCs: hematopoietic stem cells. For further details please refer to Materials and Methods.

### The NK cell score is associated with improved patient survival

It was recently shown that NK cells play a critical role in the initiation of a robust immune response against melanoma, secreting the chemokines XCL1 and CCL5 [2] and the FLT3 ligand [58] to recruit conventional dendritic cells and promote their functions. NK-cell mediated killing of tumor cells facilitates DC phagocytosis of tumor cells, and, following the migration of these cells to lymph nodes, this ultimately allows the development of a robust T-cell response against tumor antigens (summarized in Fig. 1). In agreement with this, tumors which have a high NK score are associated with much better patient survival (Fig. 4A), consistent with the results from Böttcher *et al* (2018) and a range of *in vivo* animal survival studies [5, 15, 59]. This effect is largely recapitulated by the important NK cell-secreted chemokines *XCL1* and *CCL5* (Fig. 4B & C), as well as the NK cell effector *GZMB* (Fig. 4D), and tumor-expressed *FAS* (Fig. 4E) for targeting by NK expressed *FASLG* (Fig. 3). Increased expression of NK cell adhesion protein *CD96* is also associated with improved survival (Fig. 4F).

**Figure 4:**
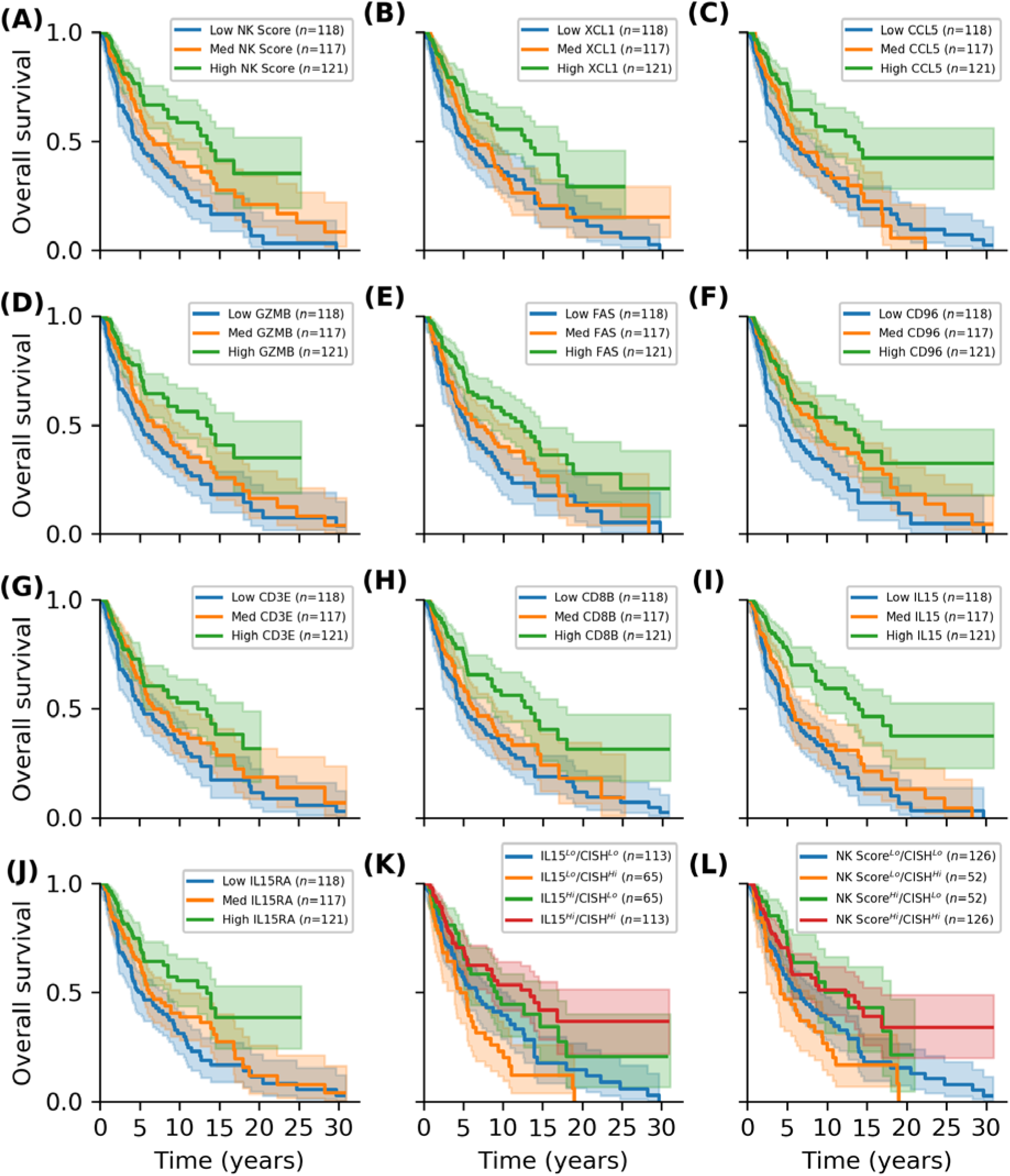
Survival outcomes for TCGA SKCM patients. Patients with metastatic tumours were partitioned by predicted NK cell infiltration as well as other indicated markers/gene set scores and associated Kaplan-Meier survival functions were examined.

We note that it is difficult to disentangle the survival effects of other cytotoxic/effector immune cells, as improved survival effects are also seen for the T cell markers CD3E and CD8B (Fig. 4G & H). This is reflected by the very strong association between our NK score, and scores derived from the TCGA Immune cluster genes (Fig. S6A) or the T cell signature (Fig. S6B), which also similar survival effects (Fig. S6C & D). As discussed above (Fig. 1), however, there is increasing evidence that NK cells play a key role initiating the intercellular signaling cascade which is necessary for strong immune recruitment. Using the Böttcher 5 gene NK cell signature [2] there is good concordance with our NK signature score (Fig. S6E), as well as between our NK score and a score calculated using the Böttcher 4 gene DC cell signature (Fig. S6F), again with similar survival effects (Fig. S6G & H).

The cytokine IL-15 has received increasing attention as a potent cytokine to increase the activity of immune cells including NK cells. Accordingly, higher transcript abundance of IL15 (Fig. 4I) as well as the receptor subunit IL15RA (Fig. 4J) are associated with improved patient survival. The gene encoded by CISH is a potent negative regulator of IL-15 induced signaling in NK cells [16] and intriguingly, tumors with high *IL15* transcript show a protective effect associated with high CISH abundance, while tumors with low IL15 showed better patient survival when CISH was low (Fig. 4K). When NK score is compared to CISH abundance we note CISH abundance has little survival effect on the high NK score subset, whereas low CISH is still associated with improved survival for patients who have tumors with a low NK score (Fig. 4L). It should be noted, however, that there is a positive association between IL15, CISH and NK score abundance, and accordingly the Lo/Lo and Hi/Hi patient subsets are larger for these comparisons. It is tempting to speculate that this association reflects activation of CISH expression in response to active IL-15 signaling, particularly within the NK high tumor subset; conversely, within the low NK cell subset, higher expression of CISH appears to be particularly deleterious.

### NK cell targeting of mesenchymal-like melanoma tumors with evidence of low TGF-β activity is associated with favorable patient survival

Recent work on features of innate anti-PD-1 resistance in melanoma found similarities with markers of MAPK inhibitor resistance [60], and as noted earlier, melanoma phenotype switching has been linked to general drug/MAPK inhibitor resistance [31]. Investigating this further, we examined the relative association between our NK score, and several phenotype-associated scores, including: a proliferative, epithelial, phenotype; an invasive, mesenchymal phenotype; and a mesenchymal phenotype where EMT has been induced specifically by TGF-β [32]. We found no association between NK score and epithelial score (Fig. 5A), however there was an association with mesenchymal score (Fig. 5B), such that less mesenchymal tumors have lower NK scores, while highly mesenchymal samples had a range of NK scores, suggesting that a subset of these tumors had higher NK cell infiltration. Intriguingly, there was no association with NK score relative to the TGF-β specific EMT gene score (Fig. 5C), despite the fact that there was a relatively strong positive association between TGF-β EMT score and mesenchymal score (Fig. 5D) – as we had previously observed however, while all samples with a high TGF-β EMT score scored high against general mesenchymal gene expression, a subset of highly mesenchymal samples showed no evidence of TGF-β driven EMT. Further, while neither the TGF-β EMT score (Fig. 5E) or NK score (Fig. 5F) had any association with age, when we partitioned patients by NK score and TGF-β EMT score, those with evidence of good NK cell infiltration and low TGF-β activity had favorable survival outcomes (Fig. 5G & H), particularly for patients within the younger cohort.

**Figure 5:**
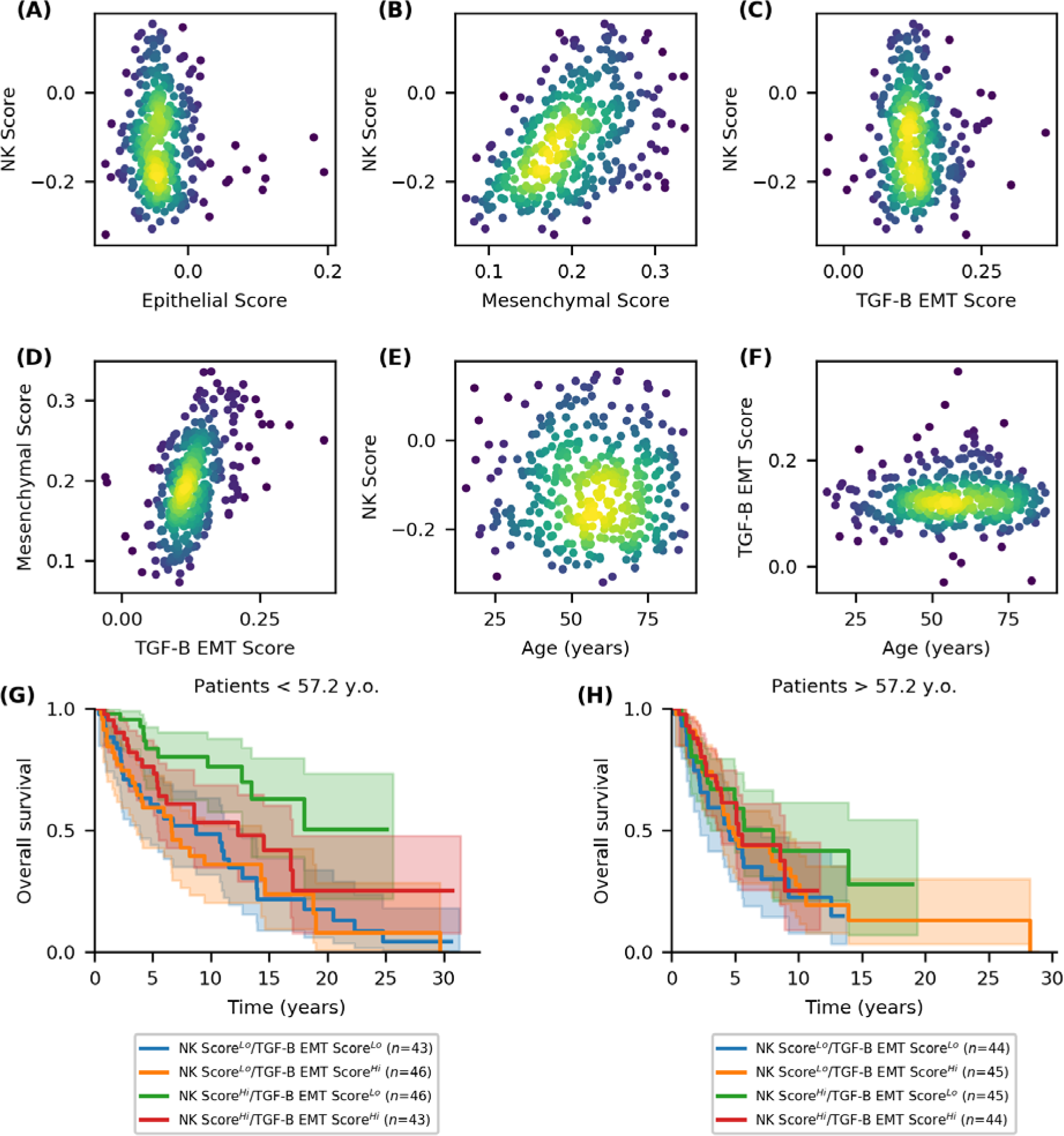
Melanoma tumours with evidence of a mesenchymal-like phenotype but low TGF-β activity, and a high NK score show favourable patient outcomes. Associations across the TCGA SKCM metastatic tumour samples, between (**A**-**C**) NK score and scores associated with EMT/phenotype-switching, (**D**) mesenchymal score and a score of specific TGF-β induced EMT; and (**E**, **F**) NK score or TGF-β EMT score and age. (**G**, **H**) Kaplan-Meier survival curves for patients partitioned by TGF-β EMT score and NK score, and split by age.

Although most *in vivo* experiments indicate a primary role for NK cells in limiting metastatic colonization [6], these data suggest that not only are NK cells associated with established metastatic tumors, but the presence of NK cell infiltrate is associated with an improved prognostic outcome.

### Natural killer cells offer a promising avenue for targeted immunotherapeutics to control melanoma metastasis

As noted above, a critical role for NK cells in driving a robust immune response has further support from a recent study which has demonstrated an important role for the NK-DC cell axis in modulating responsiveness to immunotherapy [58]. To further investigate potential modulators of NK cell infiltration we next examined transcriptomic data from the LM-MEL panel which contains representative cell lines for both the proliferative and invasive phenotype. Gene sets were filtered to retain only those present in both the TCGA and LM-MEL data, and gene set scoring was repeated for both data sets to facilitate comparison between tumor samples and the corresponding cell line models (Fig. S7A-C). As shown, in the absence of a high NK score, patients with a high mesenchymal score show no survival effects associated with the TGFβ-EMT score (Fig. S7D).

A number of melanoma cell lines from the LM-MEL panel appear to be associated with various subsets of high/low mesenchymal score and TGFβ-EMT score (Fig. S7A-C; colored scatter markers). By contrasting genes correlated or anti-correlated with NK score across the TCGA data against these cell line data we can identify markers which may be derived from the melanoma tumor which exert an immunomodulatory effect (Fig. S7E). To demonstrate the association with phenotype-switching the markers CDH1 and MITF are included [29, 34]. Further, TGF-β activity in these cell lines has a demonstrated association with *THBS1* [33]. Consistent with our observation that NK score tends to be higher in more mesenchymal tumor samples, many of the positively correlated genes tend to have higher expression in the MITF-low cell lines. Similarly, many of the anti-correlated genes tend to have lower expression in the MITF-low cell lines.

Several notable genes are present within these lists (Fig. S7E). Again, B2M is identified, together with a number of HLA-genes, suggesting that more mesenchymal-like melanomas may be more immunogenic in part because of increased antigen presentation associated with this phenotype. Given data linking IL-18 to NK cell activity [12] (Fig. 1), it is interesting to note that in the more mesenchymal cell lines, the expression of IL-18 appears to be slightly higher in the TGF-β EMT low samples. From the genes anti-correlated with NK score we note that CMTM4 was recently identified as a positive regulator of PD-L1 (together with its paralog CMTM6 which is not present) [61, 62] and this appears to have lower expression within more mesenchymal cell lines.

### Discussion

Immune “checkpoint” inhibitor antibodies, which function by reactivating tumor-resident cytotoxic lymphocytes, have revolutionized cancer therapy. Although much research is currently directed towards programs that underlie immunotherapy resistance, we lack an in-depth understanding of the fundamental mechanisms dictating response and we do not yet have robust markers to identify patients who may respond preferentially in the context of metastasis. Checkpoint inhibitors primarily block inhibitory pathways in tumor-resident T cells, however interest in other effector populations, such as NK cells, is growing [9], with recent studies showing that NK cells have a critical role in immunotherapy success [58].

Clinically, NK cell activity has been shown as inversely correlated with cancer incidence [63]. More recent evidence has shown that NK cell infiltration in human tumors is associated with better prognosis in squamous cell lung, gastric and colorectal carcinomas [6]. In melanoma cells, researchers have previously found specific HLA-I allelic losses in up to 50% of patients analyzed, and even when expressed on melanoma cells, specific HLA class I molecules are often at insufficient levels to inhibit NK cell-mediated cytotoxicity [64]. These data suggest that metastatic melanoma is an ideal target for NK cell-mediated killing and therapies that enhance NK cell activity should be investigated further. Accordingly, we have performed a detailed investigation of the transcriptomic and matched clinical data available through the TCGA skin cutaneous melanoma cohort [38].

Consistent with previous reports which have elucidate the links between dendritic cells and natural killer cells [2, 58], we show that inferred NK cell infiltration is associated with improved patient survival. We note that while the gene set derived by Böttcher and colleagues performs well [2], they calculated NK scores using mean log_2_ abundance data which can be susceptible to outliers and places a greater weighting on genes with high transcript abundance. As demonstrated (Fig. S6E & S6F), while *singscore* has been developed for larger gene sets, it still performs relatively well with the 5 gene Böttcher NK signature (NCR3, KLRB1, PRF1, CD160, NCR1). While it is hard to directly compare the accuracy of these signatures without validation data for NK cell infiltration, these results demonstrate the application of our computational method in estimating the abundance and heterogeneity of different immune subsets across different tumors and patients. Importantly, our methods allow variations in these relative immune scores to be easily compared against other relevant phenotype-associated gene sets as demonstrated by the striking survival effects we have observed when our NK score is examined in the context of melanoma phenotype switching and TGF-β signaling (Fig. 5G & H).

The observation that the inhibitory receptor CD96 appears to have a protective effect, with high CD96 corresponding to improved patient survival (Fig. 3F) is interesting since murine data has proposed that CD96 acts as a NK cell checkpoint limiting NK cell anti-melanoma immunity [65]. Furthermore, while both CD96 and TIGIT have been explored as potential immunotherapy targets [66], it has previously been shown that TIGIT appears to have a dominant role over CD96 as a checkpoint in NK activity [67]. Indeed, a recent murine report has evidenced a potent inhibitory role for TIGIT on NK cell anti-tumor immunity [68]. Further, while there are prominent long-term survival effects associated with patient age (Fig. 2B), we note that there is no association between NK score and age (Fig. 5E), yet the survival advantages associated with a low TGFβ-EMT score and a high NK score are particularly pronounced within younger patients (Fig. 5G). These results suggest that younger patients may receive a greater benefit from NK targeted immunotherapies, perhaps reflecting a higher capacity of the immune system in young patients to produce a robust anti-tumor response.

The NK cell gene signature and NK cell gene score that we describe here can be readily applied to other cancer datasets which are becoming increasingly available thanks to the efforts of large cancer research consortia. The information from such gene signature analyses will also allow researchers to stratify responders and non-responders to conventional treatments, identify the patients that are likely to profit from NK cell-based immunotherapies, and facilitate the development of prognostic markers for personalized immunotherapeutics.

## Authors’ Contributions

**Conception and design**: J. Cursons, F. Souza Fonseca Guimaraes, N.D. Huntington, A. Behren, M.J. Davis

**Development of methodology**: J. Cursons, M. Foroutan, M.J. Davis

**Analysis and interpretation of data** (e.g., statistical analysis, biostatistics, computational analysis): J. Cursons, A. Anderson, S. Hediyeh Zadeh, F. Souza Fonseca Guimaraes, N.D. Huntington, M. Foroutan, M.J. Davis

**Writing, review, and/or revision of the manuscript**: J. Cursons, F. Souza Fonseca Guimaraes, N.D. Huntington, A. Behren, M.J. Davis

**Administrative, technical, or material support** (i.e., reporting or organizing data, constructing databases): J. Cursons, A. Anderson, M. Foroutan

**Study supervision**: J. Cursons, F. Souza Fonseca Guimaraes, N.D. Huntington, M.J. Davis

## Acknowledgments

This work is supported in part by project grants from the National Health and Medical Research Council (NHMRC) of Australia (#1147528 to J.C.; #1128609 to M.J.D.; #1124784, #1066770, #1057852, #1124907 to N.D.H; and #1140406 to F.S.F.G). F.S.F.G. was supported by NHMRC Early Career Fellowship (1088703), National Breast Cancer Foundation (NBCF) Fellowship (PF-15-008), and grant #1120725 awarded through the Priority-driven Collaborative Cancer Research Scheme and funded by Cure Cancer Australia with the assistance of Cancer Australia. N.D.H is a NHMRC CDF2 Fellow (1124788), a recipient of a Melanoma Research Grant from the Harry J Lloyd Charitable Trust, Melanoma Research Alliance Young Investigator Award, and a CLIP grant from Cancer Research Institute. M.J.D was supported by NBCF Career Development Fellowship ECF-14-043 and is the recipient of the Betty Smyth Centenary Fellowship in Bioinformatics. This study was made possible through Victorian State Government Operational Infrastructure Support and Australian Government NHMRC Independent Research Institute Infrastructure Support scheme.

Results published here are based upon data generated by the TCGA Research Network: http://cancergenome.nih.gov/.

